# Myanmar’s terrestrial ecosystems: status, threats and conservation opportunities

**DOI:** 10.1101/2020.08.18.256750

**Authors:** Nicholas J. Murray, David A. Keith, Adam Duncan, Robert Tizard, Jose R. Ferrer-Paris, Thomas A. Worthington, Kate Armstrong, Nyan Hlaing, Win Thuya Htut, Aung Htat Oo, Kyaw Zay Ya, Hedley Grantham

**Author notes:** **Corresponding author:** Nicholas Murray, College of Science and Engineering, James Cook University, Townsville, Queensland, Australia. 4811.

## Abstract

Myanmar is highly biodiverse, with more than 16,000 plant, 314 mammal, 1131 bird, 293 reptile, and 139 amphibian species. Supporting this biodiversity is a variety of natural ecosystems—mostly undescribed—including tropical and subtropical forests, savannas, seasonally inundated wetlands, extensive shoreline and tidal systems, and alpine ecosystems. Although Myanmar contains some of the largest intact natural ecosystems in Southeast Asia, remaining ecosystems are under threat from accelerating land use intensification and over-exploitation. In this period of rapid change, a systematic risk assessment is urgently needed to estimate the extent and magnitude of human impacts and identify ecosystems most at risk to help guide strategic conservation action. Here we provide the first comprehensive conservation assessment of Myanmar’s natural terrestrial ecosystems using the IUCN Red List of Ecosystems categories and criteria. We identified 64 ecosystem types for the assessment, and used models of ecosystem distributions and syntheses of existing data to estimate declines in distribution, range size, and functioning of each ecosystem. We found that more than a third (36.9%) of Myanmar’s area has been converted to anthropogenic ecosystems over the last 2-3 centuries, leaving nearly half of Myanmar’s ecosystems threatened (29 of 64 ecosystems). A quarter of Myanmar’s ecosystems were identified as Data Deficient, reflecting a paucity of studies and an urgency for future research. Our results show that, with nearly two-thirds of Myanmar still covered in natural ecosystems, there is a crucial opportunity to develop a comprehensive protected area network that sufficiently represents Myanmar’s terrestrial ecosystem diversity.

## 1. Introduction

South-east Asia is an important centre of global biodiversity, with ecosystem diversity that includes tropical and temperate forests, seasonal wetlands and alpine ecosystems (Ashton & Seidler 2014). However, dense human populations, rapid economic growth and an expanding footprint of extractive activities have impacted natural environments across the region, with losses of ecosystems accelerating rapidly from around the second half of the 20^th^ century (Wilcove et al. 2013). Myanmar’s natural ecosystems provide essential ecosystem services including food, water treatment and other basic human needs to millions of people, and many have deep cultural and religious significance (Aung 2007; Barrow 2019). Despite their value, the status of Myanmar’s natural ecosystems has not been systematically evaluated, leading to uncertainty about when and where to implement conservation actions.

Myanmar is the second largest Southeast Asian country, with a terrestrial extent of about 676 600 km^2^ containing a human population of around 53.71 million (The World Bank 2019). Myanmar has very high species diversity, including more than 16 000 species of plants, and 314 mammal, 1131 bird, 293 reptile, and 139 amphibian species (Ministry of Environmental Conservation and Forestry 2014; Francis 2019; Middleton et al. 2019). The country spans a wide latitudinal range (~18.5°) and an elevational gradient ranging from sea level to over 5850 m, intersecting 19 global ecoregions (Dinerstein et al. 2017). It has large tracts of forest structured by precipitation and temperature gradients that extend across tropical evergreen lowlands and dry subtropical rain shadows to temperate mountain slopes in the eastern Himalayas. A wide central floodplain associated with the Ayeyarwady River supports monsoonal wetlands, dry forest, and savannah, while coastal areas are fringed by mangrove, tidal mudflat, sandy beach and other coastal ecosystems. Myanmar has biogeographical links to the Sundaic, East and South Asian, and Himalayan regions (Ashton & Seidler 2014) and has a considerable maritime influence with nearly 6,300 km of coastline bordering the Bay of Bengal and the Andaman Sea (Figure 1). The largest remaining contiguous patches of natural ecosystems are located as a wide band across the north and north-east in Chin, Sagaing and Kachin states, and in the south-east, including Tanintharyi and Kayin states.

**Figure 1.**
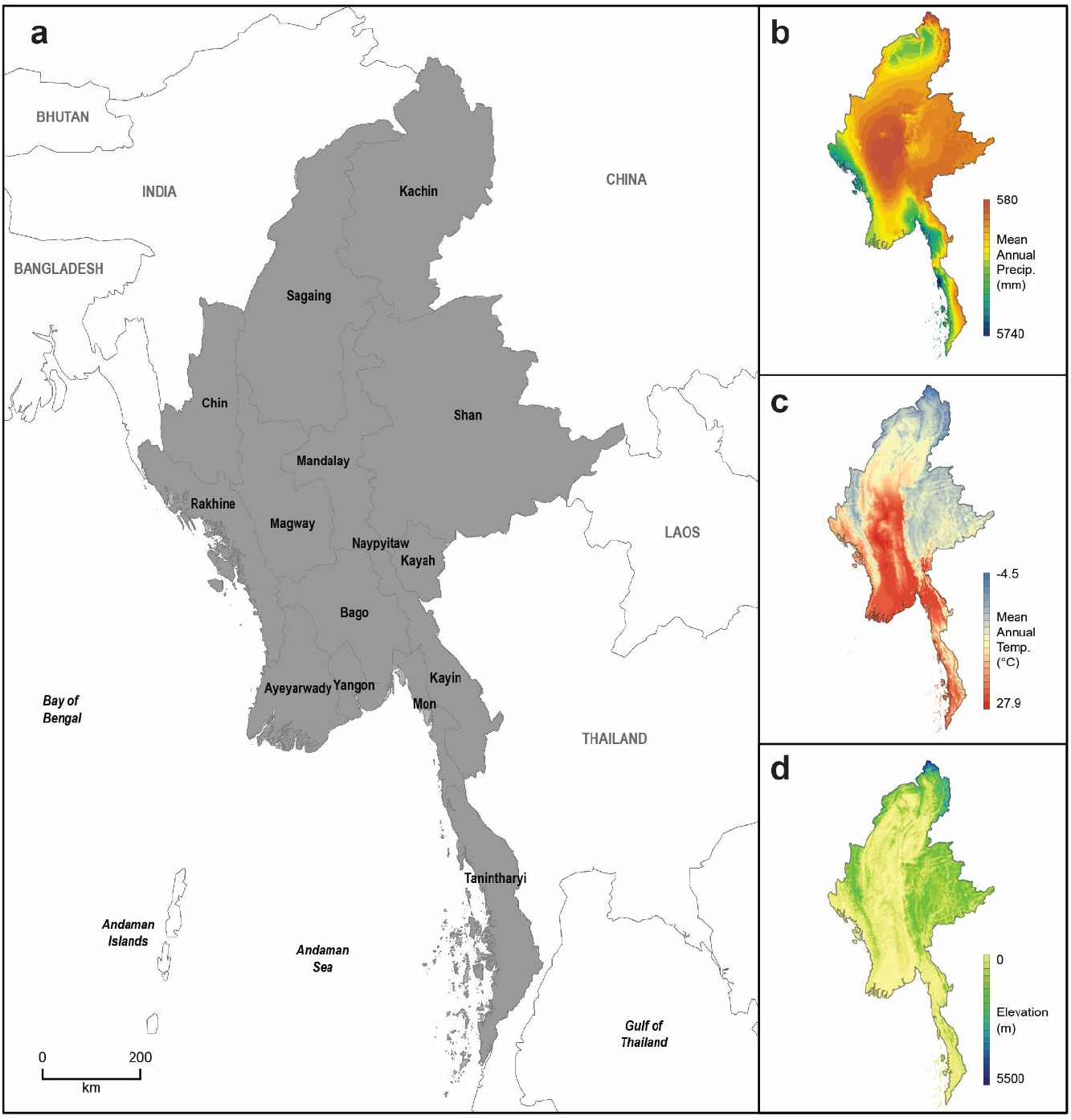
Map of Myanmar and the main climate and topographical drivers of ecological diversity. The panels show the distribution of (a) the states of Myanmar, (b) mean annual precipitation, (c) mean annual temperature, (d) elevation (data sourced from WorldClim; Hijmans et al. 2005).

The unique socio-political history of Myanmar has enabled a larger portion of these ecosystems to persist compared to neighbouring countries, although some have undergone significant recent degradation driven by rapidly intensifying threats (Lim et al. 2017; Prescott et al. 2017; Zhang et al. 2018; De Alban et al. 2020). Despite the high diversity and global significance of Myanmar’s ecosystems, they remain remarkably poorly documented. The few studies that have attempted to describe the extent of change of Myanmar’s environment have typically focused on net declines in forest extent, and have not identified ecosystems at risk of loss or priorities for protection (Webb et al. 2014; Connette et al. 2016; Bhagwat et al. 2017; De Alban et al. 2020). Owing to decades of inaccessibility, new species from Myanmar are regularly described, such as 12 new gecko species recently discovered in karst ecosystems in the Salween River Basin and Shan hills (Grismer et al. 2017). Myanmar has now become a focal region of research and conservation efforts aiming to stem losses and degradation of both species and ecosystems.

Efforts to build natural history collections have increased in recent years (Ito & Barfod 2014) and remote sensing studies of forest cover provide overviews of change for a select few ecosystem types, such as broadly defined tropical forests and mangroves (Songer et al. 2009; Webb et al. 2014; De Alban et al. 2018; De Alban et al. 2020). Local descriptive studies of ecosystems are also beginning to emerge (Oo & Koike 2015; Khaing et al. 2019). However, the primary source for a national inventory of Myanmar’s ecosystems is more than 90 years old (Stamp 1925b), with subsequent updates (Davis 1960; Kress et al. 2003) adding little new content. The lack of a systematic inventory of Myanmar’s ecosystems hampers the identification of ecosystems undergoing loss and degradation, as well as limiting the prioritization and coordination of conservation actions. At present, lists of threatened megafauna species remain the primary driver of conservation decisions (Ministry of Environmental Conservation and Forestry 2014), but these fail to represent the full range of biodiversity, ecosystem functions and services that require protection across the country (Keith et al. 2015). A systematic inventory and analysis of risks to Myanmar’s ecosystems is therefore essential for identifying more broadly based conservation priorities, designing a representative protected area network, and developing ecosystem management strategies to promote sustainable development.

The IUCN Red List of Ecosystems is the global standard for assessing the risk of ecosystem collapse, requiring systematic assessment of five criteria that focus on various indicators of ecosystem collapse (Keith et al. 2013; Rodríguez et al. 2015). The criteria require systematic analyses of change in area, range size, environmental degradation and biotic disruption over several time frames to estimate the status of each ecosystem (Rodríguez et al. 2015). National Red lists of ecosystem assessments are becoming a widely applied tool for informing environmental planning and management and for designing national protected area networks around the world (Keith et al. 2015; Alaniz et al. 2019; Bland et al. 2019).

Here we assess risks to Myanmar’s terrestrial ecosystems using the IUCN Red List of Ecosystems categories and criteria. A list of ecosystem types is requisite for a Red List of Ecosystem assessment; we used historical accounts, recent local surveys, information from experts, and extensive field reconnaissance to identify 64 terrestrial ecosystem types occurring in Myanmar. We used models of ecosystem distribution developed from remote sensing and environmental data, to assess the distribution size of each ecosystem, and synthesised existing information to estimate declines in distribution and, where possible, degradation of each ecosystem type over multiple assessment timeframes. Thus, we developed the first list of threatened ecosystems for Myanmar, and discuss the implications of our findings in the context of ecosystem conservation in Southeast Asia.

## 2. Materials and Methods

### 2.1 Study region

We included Myanmar’s entire terrestrial land mass in our analysis, including offshore islands and the intertidal zone. We conducted all of our analyses at the national-scale and summarised the data by 15 state jurisdictions (Fig. 1) to support national environmental planning.

### 2.2 Ecosystem assessment units

To develop a list of candidate terrestrial ecosystems for Myanmar we initially identified functional groups of ecosystems likely to be represented within the country by reviewing historical accounts (Stamp 1925b; Kingdon-Ward 1944; Davis 1960; Kress et al. 2003), regional reviews (e.g. Ashton & Seidler 2014) and published studies of specific ecosystem types (e.g. Oo & Lee 2007; Oo & Koike 2015; Khaing et al. 2019). For the assessment we focused on natural ecosystem types, and excluded anthropogenic ecosystem types. The IUCN global ecosystem typology (Keith et al. 2020) enabled us to structure the review of Myanmar’s ecosystems, identify similar types described in various studies, compare their descriptions, and rationalise them into a draft list of candidate ecosystems to serve as ecosystem assessment units (Rodríguez et al. 2015). It also enabled us to identify apparent gaps in previous studies, such as Myanmar’s alpine ecosystems and savannas, which are widespread but have been neglected in the majority of previous accounts and remain poorly known.

After developing a candidate list of ecosystems for Myanmar, we convened two workshops with experts from academic institutions, the Myanmar Forestry Department and local NGOs in Naypyitaw, Myanmar. Experts were asked to review the candidate list of ecosystem types, identify any gaps and, where possible, describe the spatial distribution of each ecosystem type using the state jurisdictions as a template. Following the workshops, we traversed more than 3,800 kilometres of Myanmar to investigate the distribution of ecosystem types and collect descriptive information about their composition, structural features and environmental relationships (Murray et al. 2020b). During the field traverses we also collected spatially explicit records of ecosystem occurrences for use as training data for the ecosystem mapping component of the assessment (Murray et al. 2020a).

The final list of terrestrial ecosystems for Myanmar included 64 ecosystem types from 19 ecosystem functional groups, representing 10 biomes (Table 1). For these, we compiled comprehensive ecosystem descriptions from literature reviews and our field data to summarise the characteristic abiotic and biotic elements, key ecosystem processes, distribution and proximate threats for each ecosystem. The descriptions were reviewed and, where appropriate, amended by experts, and developed as a guide to the ecosystems of Myanmar (Murray et al. 2020b).

**Table 1.** Area and protection of Myanmar’s terrestrial biomes. Not all ecosystems were mapped during remote sensing analyses and these estimates are considered minima.

| Biome | No. of ecosystems described | No. of ecosystems mapped | Mapped area of biome | Biome in relation to size of Myanmar | Biome in relation to extent of remaining ecosystems Myanmar | Extent within protected areas |  |
| --- | --- | --- | --- | --- | --- | --- | --- |
|  |  |  | km<sup>2</sup> | % | % | km<sup>2</sup> | % |
| Brackish tidal systems | 5 | 4 | 6310.9 | 0.9 | 1.5 | 133.8 | 2.1 |
| Dry subterranean | 1 | 1 | 0.4 | 0.0 | 0.0 | 0.0 | 2.6 |
| Lakes | 1 | 1 | 21.5 | 0.0 | 0.0 | 21.0 | 97.6 |
| Palustrine wetlands | 5 | 4 | 4057.3 | 0.6 | 1.0 | 0.8 | 0.0 |
| Polar/alpine | 4 | 4 | 3811.5 | 0.6 | 0.9 | 2756.6 | 72.3 |
| Savannas and grasslands | 12 | 10 | 29268.0 | 4.3 | 6.9 | 758.2 | 2.6 |
| Shoreline systems | 2 | 2 | 4458.6 | 0.7 | 1.0 | 18.8 | 0.4 |
| Supralittoral coastal systems | 2 | 0 | - | - | - | - | - |
| Temperate-boreal forests & woodlands | 7 | 6 | 46959.9 | 6.9 | 11.0 | 12208.0 | 26.0 |
| Tropical & subtropical forests | 25 | 24 | 331740.2 | 49.0 | 77.8 | 24323.4 | 7.3 |
| Total | 64 | 56 | 426628.15 | 63.1 | 100.0 | 40220.6 | 9.4 |

### 2.3 Assessing risk of ecosystem collapse

We followed the IUCN Red List of Ecosystems guidelines (Bland et al. 2017) to assess each ecosystem under the five assessment criteria that relate to indicators of the risk of ecosystem collapse. These are: reduction in geographic distribution (Criterion A); restricted geographic distribution (Criterion B); environmental degradation (Criterion C); disruption of biotic processes or interactions (Criterion D); and probability of collapse (Criterion E). We collated available published data to enable assessments of all of the criteria for which information was available. The application of the red list criteria resulted in each ecosystem being assigned a status based on the most threatened outcome of any of the criteria and sub-criteria included the bounds of uncertainty in the status of each ecosystem (Bland et al. 2017). If a criterion could not be assessed due to lack of data it was denoted *Data Deficient* or, when data or a method was available but not able to be used in the assessment, as *Not Evaluated* (Bland et al. 2017).

#### 2.3.1 Ecosystem geographic distribution and change data

We assessed changes in ecosystem extent (Criterion A) using publicly available spatial data (e.g., Murray et al. 2019; Murray et al. 2020a) and published estimates of area change (e.g., Webb et al. 2014). As the data were available for timeframes varying from 18 to 27 years, we extrapolated area change estimates to the 50-year time frames using the exponential decline function from the R package *redlistr* (Lee et al. 2019). Our satellite analyses of ecosystem distributions provided the basis for estimating range size (Criterion B) and for delineating spatial boundaries for analysing ecosystem degradation (Murray et al. 2020a; Murray et al. 2020b). Owing to uncertainty in the distribution of some ecosystems, only 57 (89.1%) of Myanmar’s ecosystem types could be included in mapping analyses. We developed a Google Earth Engine module to compute Area of Occupancy and Extent of Occurrence of each ecosystem (Criterion B), and inferred continuing declines and threats from published studies and expert elicitation.

#### 2.3.2 Environmental degradation

We used results from existing studies (n = 3 ecosystem types) and environmental suitability models (n = 33 types) to assess future environmental degradation (Criterion C) due to climate change. For three mangrove ecosystems, we used data from Lovelock et al. (2015) to estimate the proportion of each ecosystem likely to be drowned by 2060 under three sea-level rise scenarios (0.48m, 0.63m and 1.4m of sea level rise by 2100).

For ecosystems amenable to bioclimatic modelling (n = 33) we developed environmental suitability models to assess projected changes in bioclimatic suitability (Criterion C; Ferrer-Paris et al. 2019). We used training data from the remote sensing analyses (n = 57 955 points; Murray et al. 2020a) to identify existing environmental conditions for each ecosystem type using 19 standard bioclimatic variables as covariates in a random forest classification model (WorldClim v1.4; 1960-1990; Hijmans et al. 2005). We removed training points of the same class that fell within the same pixel of the bioclimatic variables to remove duplicate observations, and randomly partitioned remaining data into spatially stratified training (75%) and testing (25%) subsets. We set the number of covariates per decision-tree to six, the number of trees to 2000, and used stratified random sampling (by ecosystem type) of training data in each fitted tree to allow balanced representation of all ecosystems. We discarded results where classification error was >20% for both training the testing samples and where area under the sensitivity and specificity curves (AUC) for the focal class was less than 85%. The predicted suitability for each ecosystem was calculated as the proportion of 2000 classification trees assigning the ecosystem to a raster cell under expected future bioclimatic conditions according to four alternative global circulation models and four representative emission scenarios for the year 2050.

To assess the Criterion C category thresholds, suitability predictions were intersected with the extant mapped distribution of each ecosystem and the average relative severity of change in environmental suitability was estimated for each of the 16 combinations of models and emission scenarios and assessed against category thresholds over a 50 year period (2000-2050; following Ferrer-Paris et al. 2019). The collapse threshold for environmental suitability was set to the threshold of equal sensitivity and specificity identified by the bioclimatic model (see Ferrer-Paris et al. 2019). The ecosystem status under Criterion C was thus assigned the median of the 16 assessment outcomes, with plausible bounds including 90% of all outcomes.

Ecosystems with extreme uncertainty in the status outcome (plausible bounds ranging from Least Concern to Critically Endangered) were as assigned as Data Deficient to reflect severe model uncertainty (n = 10).

#### 2.3.3 Biotic disruption

To assess biotic disruption (Criterion D), we analysed the extent of ecosystem degradation with a variety of spatially explicit datasets developed from remote sensing. For the three widely distributed mangrove ecosystems (Rakhine, Ayeyarwady Delta, Tanintharyi; Supplementary Table 1), we used a newly developed 30-m remote sensing dataset that classifies pixels according to their vegetation dynamics over an 18 year period (2000-2018; Worthington & Spalding 2018). Twelve remote sensing indices relating to greenness and foliage moisture were used to identify pixels that have undergone sustained and large (>40%) decreases in greenness relative to a pre-2000 reference state (Worthington & Spalding 2018). Sustained declines in index values were assumed to be indicative of degradation, defoliation or death of mangrove trees due to the effects of anthropogenic, biotic and abiotic change (see Worthington & Spalding 2018). For each ecosystem, we estimated the proportion of the extent of the ecosystem’s distribution identified as degraded in 2000 and 2018. We linearly extrapolated these estimates to 2050 using the *redlistr* (Lee et al. 2019), assuming constant ongoing rates of exponential change to enable an assessment of mangrove degradation over a 50-year frame (Criterion D2b). Given that elements of the ecosystem remain where mangroves were mapped as degraded, we assumed the relative severity of the decline in biotic function was greater than 50% but less than 80% (Murray et al. 2020b). To estimate plausible bounds of the assessment outcome and reflect uncertainty in collapse thresholds (Bland et al. 2018), we conducted a sensitivity analysis by repeating the analysis with a relative severity of the decline assumed as >80% (Bland et al. 2017; Bland et al. 2018).

For non-mangrove forest ecosystems, we used a recently developed time-series dataset of the distribution of primary forest in Southeast Asia (Potapov et al. 2019). We computed the proportion of each ecosystem mapped as primary forest in each year of the dataset (2000-2018), and extrapolated the estimate to a 50 year time-frame using a linear model centred on the year 2009 (1984-2034; after Murray et al. 2015). We assumed the entire extent of the ecosystem was primary forest in the year 1750 and (conservatively) that ecosystem collapse occurs when the area of primary forest in an ecosystem declines to zero. The resulting estimates of change in primary forest were assessed against category thresholds (Criterion D2b), and plausible bounds of the status outcome were identified using the upper and lower confidence intervals from the linear model.

For ecosystems where primary forest data were not suitable (primarily those with estimated canopy cover < 25%; Potapov et al. 2019), we used a newly developed index representing pressure on forests and lost connectivity to assess the extent and severity of degradation since 1750 (Criterion D3; Grantham et al. 2020). The index integrates maps of changes in forest connectivity with data on human pressure known to result in ecosystem degradation to compute a continuous value of forest integrity at high resolution (ranging 0-10 for each 300-m pixel). We clipped the index data to the distribution of each ecosystem, and estimated the proportion of the ecosystem mapped with index scores in each of the following bins (0-1: > 80% relative severity; 1-3: >=70 | < 80% relative severity; 3-6: >= 50 | < 70% relative severity; >6: <50% relative severity). Bin values were set with reference to areas visited during the field trips. To represent uncertainty in these bin values we conducted a sensitivity analysis with thresholds ±0.5, which we used to set plausible bounds around the assessment outcomes.

### 2.4 Protected area coverage

We assessed protected area coverage by intersecting the ecosystem map (Murray et al. 2020a) with a curated database of Myanmar’s protected areas (Government of Myanmar, unpublished data). We also developed state-wise summaries of ecosystem diversity and status, and summarised assessment outcomes as proportions of total (Myanmar) and as proportions of each Biome.

## 3. Results

### 3.1 Status of ecosystems in Myanmar

Myanmar’s remaining natural ecosystems cover 426 628 km^2^ of the country’s land mass (Table 1; Figure 2a). On this basis, 36.9% of Myanmar’s original ecosystems have been transformed by human activities and 63.1% remains in a natural or semi-natural state (Table 1). Ecosystems in the tropical and subtropical forest biome were by far the largest group of ecosystem types identified in Myanmar (25 ecosystem types; Table 1), consisting of 14 dry forest ecosystem types, 10 lowland rainforest ecosystem types and one moist montane rainforest. This biome alone covers nearly 50% of Myanmar’s land mass and accounts for almost 80% of remaining ecosystem extent (Table 1). Savannas and grasslands were also strongly represented (12 ecosystem types), occurring in seasonally dry areas (Figure 3a), but now cover only 4.3% of Myanmar, accounting for 6.9% of remaining ecosystem extent. Temperate and subalpine forests & woodlands, consisting of 6 ecosystem types located in higher altitude areas of northern and eastern Myanmar, cover 7% of Myanmar and account for 11% of remaining ecosystem extent. Thus, tropical and subtropical forests, savannas and grasslands, and temperate and subalpine forests and woodlands account for more than 95% of Myanmar’s remaining natural ecosystem cover, and more than 60% of Myanmar’s land mass. Overall, ecosystem richness is highest in areas of high topographic and climatic diversity, primarily in Northern, Eastern and Southern Myanmar (Figure 3a; Supplementary Figure 1).

**Figure 2.**
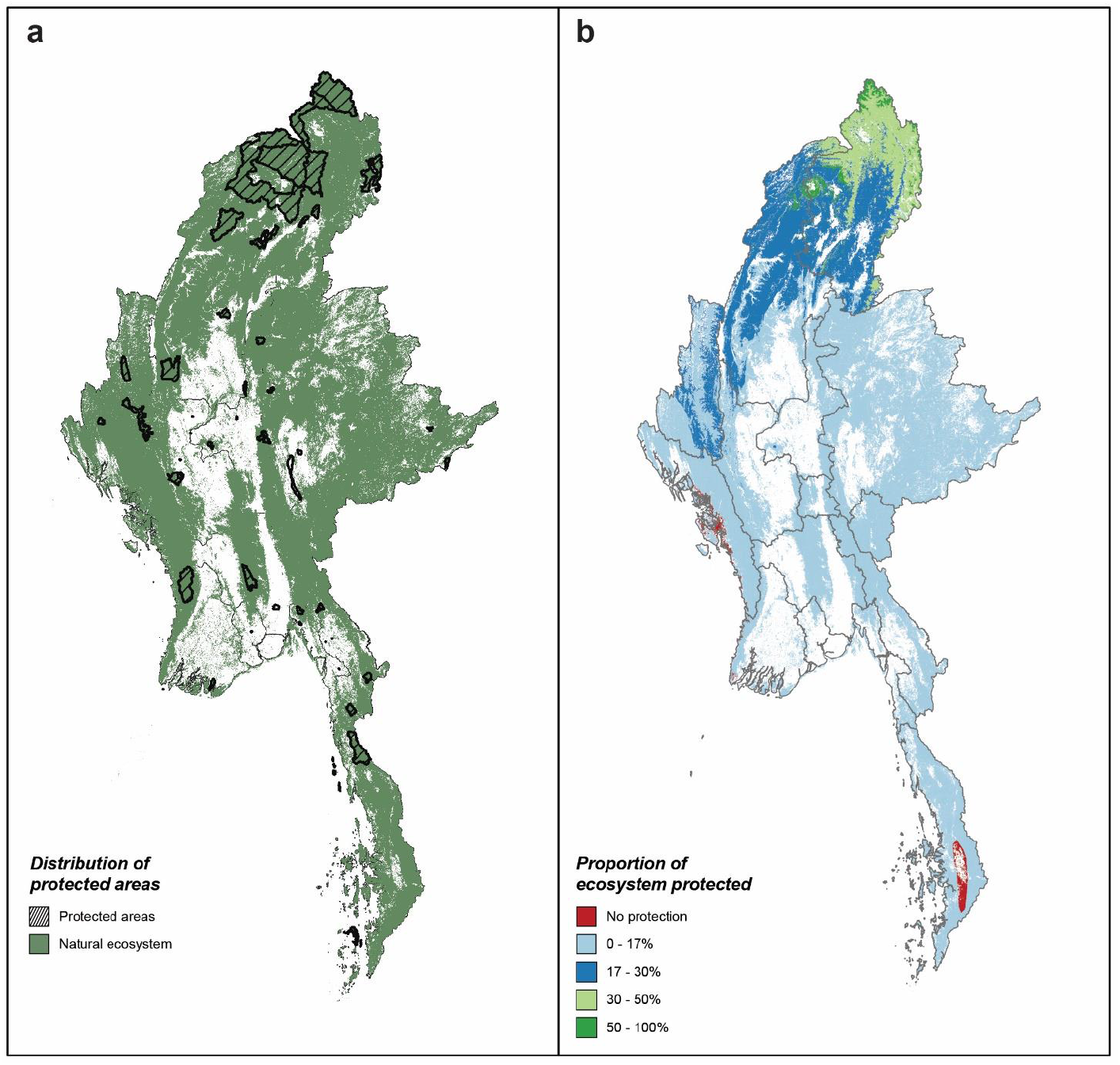
The distribution of remaining protected natural ecosystems in Myanmar. The panels show (a) the distribution of protected areas in relation to natural ecosystems and (b) the proportion of each ecosystem occurring within a protected area. Ecosystem data is from a country-wide remote sensing analysis of 57 ecosystem types, which are in varying states of degradation. Protected area data was collated by the Myanmar Government.

**Figure 3.**
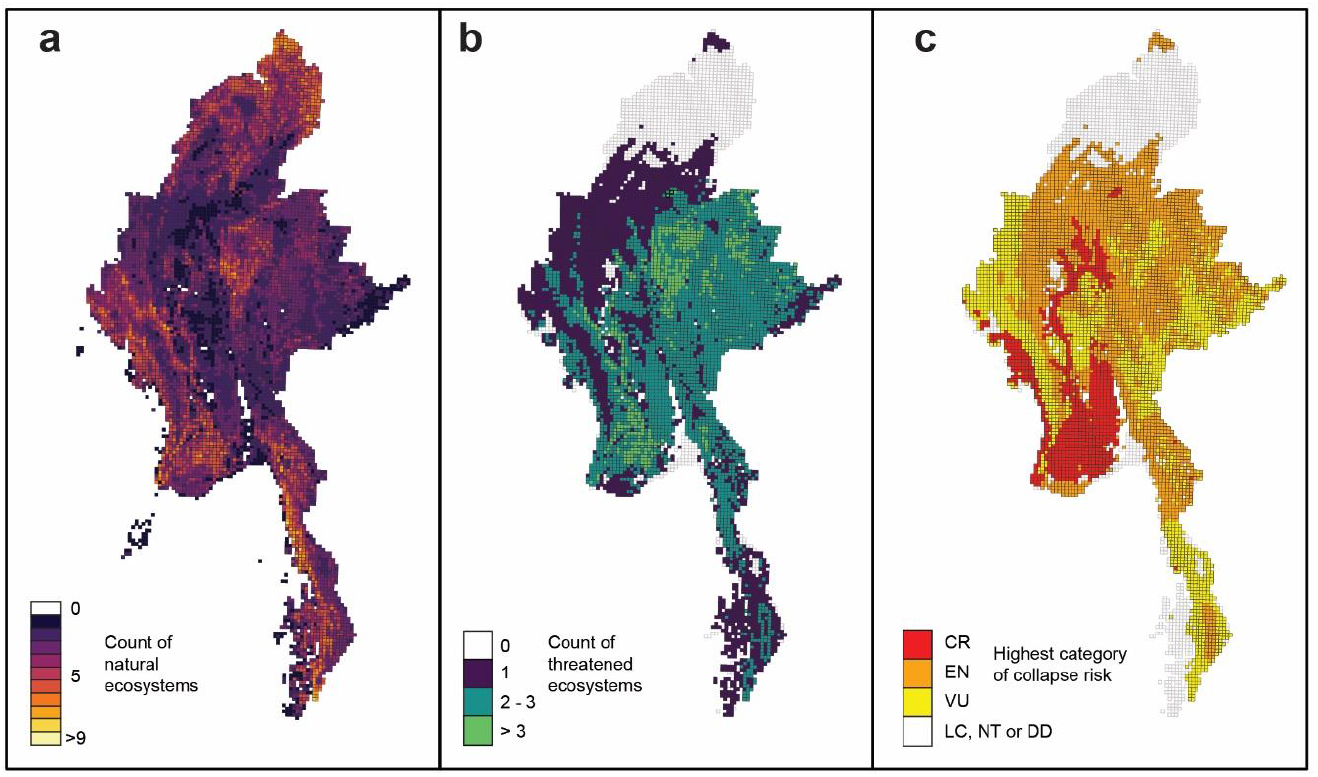
Map of the distribution of (a) all mapped ecosystems, (b) threatened ecosystems and (c) the highest risk category of ecosystem in Myanmar. The figure shows “ecosystem richness”, which represents the number of (a) ecosystems or (b) ecosystems listed as Vulnerable, Endangered or Critically Endangered under the IUCN Red List of Ecosystems that occurs in each 10 × 10 km grid cells across the country. Panel (c) the highest risk category (based on IUCN Red List of Ecosystems criteria) for any ecosystem that occurs within 10 × 10 km cells.

Twenty-nine of Myanmar’s 64 ecosystem types qualified for threatened status based on IUCN Red List criteria (Vulnerable, Endangered or Critically Endangered), while 17 were designated Data Deficient (Table 2). Approximately half of the remaining extent of natural ecosystems in Myanmar (246 478 km^2^; 57.8%) is occupied by threatened ecosystems. One ecosystem type, *Central Ayeyarwady palm savannah*, was assessed as Collapsed (Table 2) and now only seems to remain as relictual or regrowth trees with exotic ground layer plants and domestic animals in agricultural landscapes of the central Ayeyarwady basin. A further two ecosystem types assessed as Critically Endangered (CR), *Ayeyarwady kanazo swamp forest* and *Southern Rakhine evergreen rainforest*, could plausibly be Collapsed (CO), given uncertainties in mapping their distribution and no recent recorded occurrences (Table 3; Figure A.1).

**Table 2.** Summary statistics of Myanmar ecosystems assessed under the IUCN Red List of Ecosystems criteria. The number of ecosystems in each collapse risk category is listed with % threatened calculated as the number of ecosystems in the Critically Endangered, Endangered or Vulnerable categories. The highest category of risk for each ecosystem is used to assign the overall status (the Outcome Criterion).

| Biome | No. of ecosystem types | IUCN Assessment Outcome |  |  |  |  |  |  | Threatened |  | Data Deficient % | Outcome Criterion |  |  |  |  |
| --- | --- | --- | --- | --- | --- | --- | --- | --- | --- | --- | --- | --- | --- | --- | --- | --- |
|  |  | CO | CR | EN | VU | NT | LC | DD | No. | % |  | A | B | C | D | E |
| Brackish tidal systems | 5 | 0 | 2 | 1 | 0 | 1 | 0 | 1 | 3 | 60 | 20 | 2 | 1 | 0 | 0 | 0 |
| Dry subterranean | 1 | 0 | 0 | 0 | 0 | 0 | 1 | 0 | 0 | 0 | 0 | 0 | 0 | 0 | 0 | 0 |
| Lakes | 1 | 0 | 0 | 0 | 0 | 0 | 1 | 0 | 0 | 0 | 0 | 0 | 0 | 0 | 0 | 0 |
| Palustrine wetlands | 5 | 0 | 3 | 1 | 0 | 0 | 1 | 0 | 4 | 80 | 0 | 4 | 0 | 0 | 0 | 0 |
| Polar/alpine | 4 | 0 | 0 | 1 | 0 | 1 | 2 | 0 | 1 | 25 | 0 | 0 | 1 | 0 | 0 | 0 |
| Savannas and grasslands | 12 | 1 | 0 | 1 | 3 | 1 | 4 | 2 | 4 | 33.3 | 16.7 | 0 | 3 | 0 | 1 | 0 |
| Shoreline systems | 2 | 0 | 0 | 0 | 0 | 0 | 2 | 0 | 0 | 0 | 0 | 0 | 0 | 0 | 0 | 0 |
| Supralittoral coastal systems | 2 | 0 | 0 | 0 | 0 | 0 | 0 | 2 | 0 | 0 | 100 | 0 | 0 | 0 | 0 | 0 |
| Temperate-boreal forests & woodlands | 7 | 0 | 0 | 1 | 1 | 0 | 1 | 4 | 2 | 28.6 | 57.1 | 0 | 0 | 0 | 2 | 0 |
| Tropical & subtropical forests | 25 | 0 | 3 | 4 | 8 | 0 | 2 | 8 | 15 | 60 | 32 | 1 | 3 | 4 | 8 | 0 |
| Total | 64 | 1 | 8 | 9 | 12 | 3 | 14 | 17 | 29 | 45.3 | 26.6 | 7 | 8 | 4 | 11 | 0 |

CO – Collapsed; CR – Critically Endangered; EN – Endangered; VU – Vulnerable; NT – Near Threatened; LC – Least Concern; DD – Data Deficient. Note: The IUCN assessment outcome for one ecosystem (Magway dry cycad forest) was assigned from the equal outcome of assessments of Criterion C and D.

**Table 3.** Number of ecosystems in each IUCN Red List of Ecosystems conservation status category. Lower bound and upper bound reflect uncertainty in assessments of the criteria.

| IUCN Category | Overall outcome |  | Lower bound |  | Upper bound |  |
| --- | --- | --- | --- | --- | --- | --- |
|  | No. of Ecosystems | % of Ecosystems | No. of Ecosystems | % of Ecosystems | No. of Ecosystems | % of Ecosystems |
| Collapsed | 1 | 1.6 | 1 | 1.6 | 3 | 4.7 |
| Critically Endangered | 8 | 12.5 | 6 | 9.4 | 7 | 10.9 |
| Endangered | 9 | 14.1 | 10 | 15.6 | 9 | 14.1 |
| Vulnerable | 12 | 18.8 | 11 | 17.2 | 16 | 25.0 |
| Near Threatened | 3 | 4.7 | 4 | 6.3 | 2 | 3.1 |
| Least Concern | 14 | 21.9 | 15 | 23.4 | 10 | 15.6 |
| Data Deficient | 17 | 26.6 | 17 | 26.6 | 17 | 26.6 |

Of the 29 ecosystems classified as threatened, 12 were assigned to the Vulnerable category, 9 to Endangered and 8 to the Critically Endangered category (Table 2). The criteria that underpin threatened status of most of these ecosystems types indicate risks are chiefly attributable to decline in ecosystem function (15 ecosystems), from biotic degradation related to loss of primary forest or tree die-off (Criterion D; 11 ecosystems), or from abiotic changes through diminishing climatic suitability (Criterion C; 4 ecosystems). Eight threatened ecosystem types had restricted distributions susceptible to stochastic threats (Criterion B) and seven types were at risk from rapid declines in extent (Criterion A). One ecosystem, *Magway dry cycad forest*, was assigned an equal outcome from two criteria (Endangered, Criteria C2a and D3).

Three biomes, including Brackish tidal systems (60%), Palustrine wetlands (80%) and Tropical and subtropical forests (60%), had more than half of their constituent ecosystems identified as threatened. Geographically, the highest concentrations of threatened ecosystems occur in areas associated with a history of widespread conversion to agriculture and other intensive land-uses (Figure 3b). Ecosystems at imminent risk of collapse (Critically Endangered) broadly occur in areas of high human population density or highly intensive forms of agriculture such as rice cultivation (Figure 3c).

Seventeen of Myanmar’s 64 ecosystem types were classified as Data Deficient (Table 3; Table A.1). These ecosystem types are known in Myanmar from historical records or expert advice, but there was insufficient published information to assess the criteria (e.g. rocky Tanintharyi karst), or too few distribution records to incorporate into the ecosystem mapping workflow (e.g. grassy saltmarsh). Eleven of these ecosystems were initially assessed as Least Concern, but an expert review indicated there was sufficient uncertainty around this outcome to classify them as Data Deficient. Across all ecosystems, there were sufficient data to assess up to 10 of the 16 subcriteria within the five red list criteria (mean 4.5 subcriteria assessed per ecosystem type), with only three ecosystems assessed by eight or more subcriteria (three of the mangrove ecosystem types). The percentage of all subcriteria assigned to the Data Deficient category, averaged across all ecosystem types, was 66% (range 31-100%). Data from the ecosystem distribution map were sufficient to apply Criterion B in assessments of 56 of the 64 ecosystem types. Data deficiency was highest for D1 and D2a, which could not be applied to any ecosystem type, followed by C3 (97% Data Deficient), C1 (94% Data Deficient), C2B (92% Data Deficient) and A2a (91% Data Deficient).

### 3.2 Protected areas

The total protected area in Myanmar is approximately 43 538 km^2^, or about 6.4% of the country, including some 4000 km^2^ that protect assets in non-natural areas (Table 4). Protected areas cover ~9.4% (40 220 km^2^) of the remaining extent of natural ecosystems in Myanmar (Table 1), but they protect only 1.9% of the extent of threatened ecosystems. In contrast, more than 17.5% of the extent of non-threatened ecosystems are covered by protected areas, primarily in northern Myanmar. *Glacial lakes*, (21.5km^2^ total extent) are nearly entirely covered by protected areas (97.6%). The 4 ecosystem types in the polar/alpine biome are also well represented in Myanmar’s protected areas, with their 72.3% of their total extent of 3812 km^2^ occurring within large protected areas in far northern Myanmar (such as Hkakaborazi National Park and Hponkanrazi Wildlife Sanctuary; Figure 2b). In contrast, only 0.02% of the area of palustrine wetlands, which occur mainly in central Myanmar, occur within protected areas. Three ecosystem types had no protected area coverage; *Southern Rakhine evergreen rainforest, dwarf mangrove (shrubland) on shingle* and *Rakhine mangrove forest on mud* (Figure 2b; Table A.1).

**Table 4.**
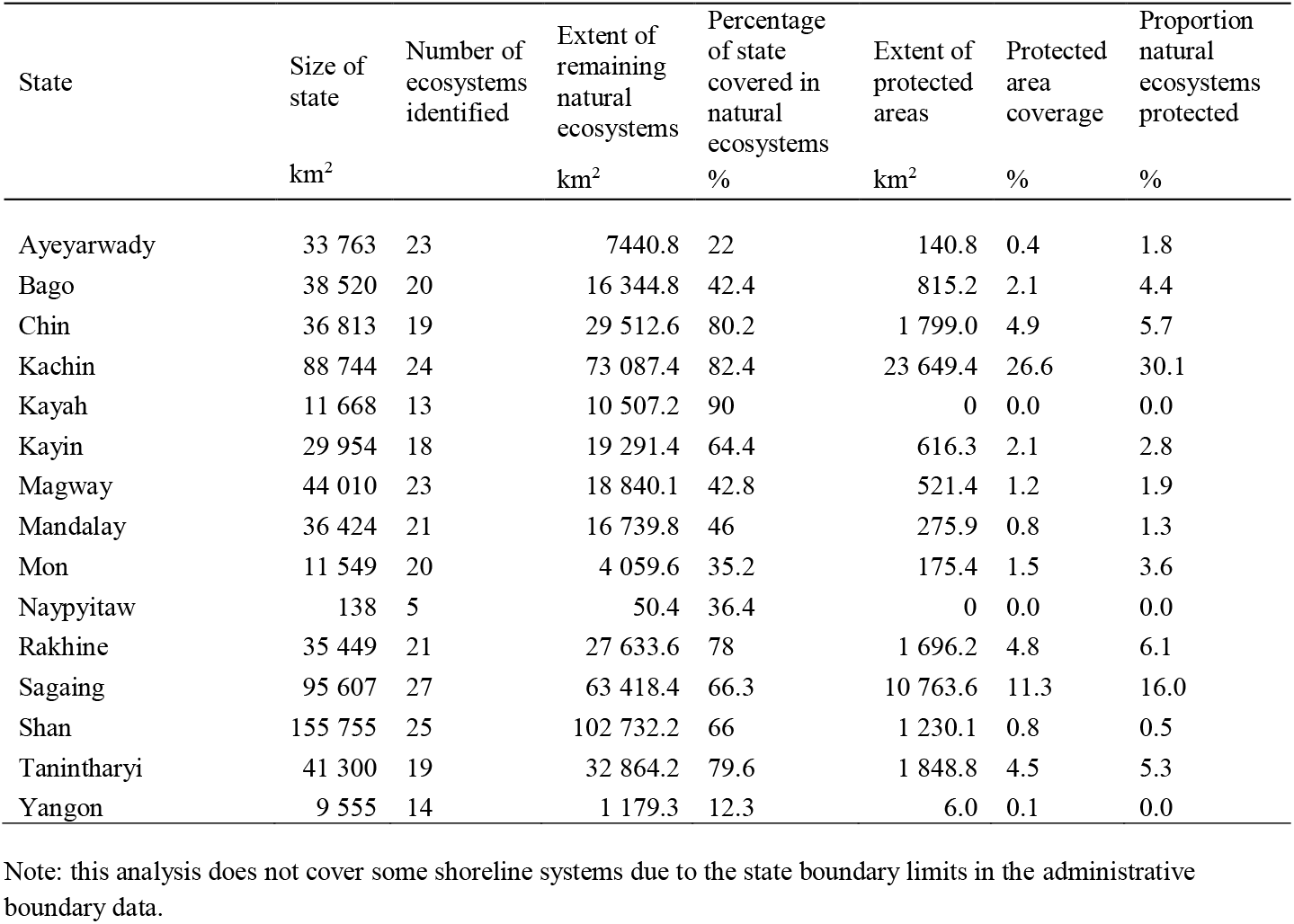
Protected areas and extent of natural ecosystems per state, Myanmar. Protected area data are sourced from a curated dataset of protected area distributions provided by the Myanmar Government.

By state, Kachin had the highest coverage of protected areas, with 23 649 km^2^ covering 30.1% of the state and protecting about 26.6% of its remaining natural ecosystems (Table 4). States with less than 1% coverage by protected areas were Naypyitaw (0%), Kayah (0%), Yangon (0.1%), Ayeyarwady (0.4%), Shan (0.8%) and Mandalay (0.8%). States with the highest percentage of natural ecosystems remaining were Kayah (90.1%), Kachin (82.4%), and Chin (80.2%), mostly consisting of ecosystem types from tropical/subtropical forests biomes (Kayah and Chin) and tropical/subtropical forest, temperate-boreal forests and woodlands, and polar/alpine biomes (Kachin). In contrast, the majority of natural ecosystems in Yangon and Ayeyarwady states have been converted to anthropogenic land uses, with natural ecosystem cover reduced to only 12.3% and 22% of each state, respectively. Sagaing had the highest ecosystem diversity (27 ecosystems) followed by Shan (25) and Kachin (24), and Naypyitaw had the fewest ecosystem types (5).

## 4. Discussion

Our assessment shows that although many of Myanmar’s ecosystems, including floodplains, lowland evergreen forests, and savannas, have undergone extensive degradation and loss, others remain as some of the most important intact examples of their kind in Asia, such as pine savannas (Ratnam et al. 2016), seasonally dry forests (Songer et al. 2009) and tropical lowland rainforests (Ashton & Seidler 2014; Connette et al. 2016). Nevertheless, we found that the majority of ecosystems in Myanmar are at risk from accelerating threatening processes, including infrastructure development, mining, tourism expansion, timber extraction, establishment of plantations for commodities such as rubber and palm oil, agricultural development, cement production, climate change and urban expansion (Leimgruber et al. 2005; Songer et al. 2009; Bhagwat et al. 2017; Hughes 2017; Lim et al. 2017). This assessment starts to fill major knowledge gaps about the diversity and distribution of ecosystems, and extent that these threatening processes are impacting ecosystems in Myanmar.

### 4.1 How much of Myanmar’s natural ecosystems remain?

Our study revealed that around 36.9% of Myanmar’s natural ecosystems have been converted to anthropogenic ecosystems over the last 2-3 centuries, and nearly half (45.3%) of Myanmar’s ecosystems now have an appreciable risk of collapse. Ecosystems at high risk of collapse show a strong spatial association with areas where crop agriculture is the dominant land use. For example, 50% of Myanmar’s Critically Endangered ecosystems and 44% of Endangered ecosystems occur in the heavily cropped central dry zone and southern Ayeyarwady floodplain, (Figure 3; Table A.1). This region has been radically transformed from natural to anthropogenic ecosystems over the last few centuries, which has been accompanied by (i) massive changes of natural water flows and inundation dynamics to support rice cultivation (Torbick et al. 2017), (ii) extensive land-clearing to support various agricultural land-uses (including peanuts, rice, aquaculture), and (iii) heavy grazing that has caused severe erosion and land degradation. The majority of palustrine wetlands, tropical dry forests and savanna ecosystems that once occurred in this region are now heavily fragmented, degraded, and remain only as very small and often degraded remnant patches (Stott 1984; Songer et al. 2009; Wohlfart et al. 2014; Ratnam et al. 2016). Other documented threats to Myanmar’s ecosystems include infrastructure development (Lim et al. 2017), logging for high value timber (Prescott et al. 2017), agricultural development (Zhang et al. 2018), plantations (Connette et al. 2016; Poortinga et al. 2019), and extraction of timber (Connette et al. 2016). Similarly, the impacts of mining for jade, tin, coal, amber, limestone and gold, particularly on Karst and forest ecosystems, have been documented in several recent studies (Bhagwat et al. 2017; Lim et al. 2017; Shimizu et al. 2017; Lee et al. 2020).

In this assessment declines in distribution (Criterion A) or risks of catastrophic threats associated with restricted distributions (Criterion B; Murray et al. 2017) determined the overall status of 50% of threatened ecosystems. The remainder were assessed as threatened due to the impacts of proximate and distal threats that have or are expected to significantly influence their abiotic or biotic function (Criteria C and D). These threats include slash and burn agriculture, high-value timber extraction, cutting for fuelwood, defaunation and climate warming (Murray et al. 2020b). Although some threats have been operating for decades (e.g. high value timber extraction) and sometimes centuries (e.g. cutting for fuelwood), others are ‘emerging’ as a result of the country’s recent economic development (e.g. new roads and oil pipelines; Rao et al. 2002). ‘Downstream’ impacts of these threats typically include degradation beyond the footprint of the impact itself, and are therefore particularly hard to identify and quantify in ecosystem assessments. Our climate projections also indicated that many of Myanmar’s ecosystems are threatened due to climate warming, and further investigations of the influence of climate change on the extent and functioning of Myanmar’s ecosystems are warranted.

During the assessment, we identified one case of ecosystem collapse in Myanmar (*Central Ayeyarwady palm savanna*). Agricultural expansion and growing regional populations in the central dry zone over the last century led to widespread degradation and conversion of this ecosystem type. Several processes likely contributed to the decline: introduced plant species outcompeted the native grassy understory. Native megafaunal engineers were extirpated, and intensive livestock grazing, in turn, limited recruitment of native species and altered natural fire regimes that maintained the structure and functioning of this savanna ecosystem. Remaining depauperate fragments of this ecosystem are few, and include scattered occurrences of former canopy species *Borassus flabellifer* that remain in the landscape after collapse has occurred. Occurrences of these species suggest palm savanna was widespread, but primarily restricted to the flat, low lying parts of the central dry zone, an area that undergoes periodic saturation associated with the monsoon and long-spells of hot dry weather that often last more than 6 months. This unique ecosystem probably once supported endemic and near-endemic birds such as Burmese Collared-dove *Streptopelia xanthocycla*, Burmese Bushlark *Mirafra microptera*, Burmese Prinia *Prinia cooki*, Ayeyarwady Bulbul *Pycnonotus blanfordi* and White-throated Babbler *Chatarrhaea gularis*, as well as a diverse assemblage of large herbivores and their predators, including tiger *Panthera tigris*. Although we found no remaining patches of this ecosystem type during field trips and inspections of high resolution satellite imagery, exhaustive targeted field searches for this ecosystem were not conducted and it is possible that a few small remnant patches remain. We therefore recommend continued investigations in the central dry zone to confirm our assessment.

Assessment results of a further two ecosystems listed as Critically Endangered could also plausibly be Collapsed. These ecosystems require urgent further field work to resolve their status (*Ayeyarwady kanazo swamp forest* and *Southern Rakhine evergreen rainforest*). The decline of the kanazo swamp forest, found at marginally higher coastal elevations than the strictly intertidal *Ayeyarawady delta mangrove forest*, began more than 100 years ago with intensive exploitation of the characteristic tree species, Kanazo (*Heritiera fames*) (Stamp 1925a). Kanazo was highly valued for construction timber, crucial during the construction of Yangon, and was also extensively cut for fuel wood (Bryant 1996). At the same time, this ecosystem underwent extensive clearing for the establishment of vast areas of rice agriculture, which now covers the majority of the lower Ayeyarwady floodplain (Stamp 1925a; Stamp 1925b; Webb et al. 2014). Even with the establishment of delta forest reserves around the turn of the century (Stamp 1925a), illegal extraction of Kanazo continued to reduce the extent of this ecosystem, with one report suggesting at least 250 000 tons of kanazo was extracted in 1919-1920 alone (Bryant 1996). Despite data searches, sufficient evidence to confirm the continued existence of this ecosystem was not found during this assessment. It is imperative that any remaining tracts of this ecosystem are identified and protected as a matter of urgency. The second Myanmar ecosystem at imminent risk of collapse is *Southern Rakhine evergreen rainforest* which, according to historical descriptions, once occurred in consistently high rainfall areas of the southern Rakhine range (Stamp 1925b). During the assessment we could not confirm its occurrence *in-situ*, but a remote sensing classification model trained from nearby evergreen rainforests in Tanintharyi state suggested that some very small patches of evergreen forest may remain within its reported range (Murray et al. 2020a). We therefore recommend an urgent field expedition to search for this ecosystem, and implement rapid conservation actions if it is confirmed in this region.

### 4.2 Addressing data deficiency

Data deficiency has been a barrier to biodiversity conservation in many countries. Myanmar is exemplary of major limitations on availability and quality of ecosystem data, including the lack of a basic ecosystem typology, country-wide maps of much of Myanmar’s biodiversity, and time-series data about ecosystem change. Despite this, our study shows that it is possible to synthesise relevant data and strategically fill gaps with new data and analyses that enabled important inferences about the status of Myanmar’s ecosystems. With the engagement of government authorities, this is proving to be instrumental for supporting conservation planning and sustainable development, and to form an agenda for future data collection. Similar benefits may be realised in other countries through productive partnerships between NGOs, academic institutions and governments.

More than a quarter (26.6%) of Myanmar’s ecosystem types qualified as Data Deficient, hindering our understanding of the status of a considerable proportion of Myanmar’s natural ecosystems. The high number of subcriteria that could not be assessed (67%) reveals a lack of data on change in ecosystem area and on the extent of degradation for the majority of ecosystems assessed. Some of our assessments relied on global (e.g. Worthington & Spalding 2018) or regional datasets (e.g. Lovelock et al. 2015; Potapov et al. 2019) and could be improved with higher resolution studies conducted at finer scales. This is, however, primarily a result of a low level of ecological monitoring over the past five decades, a lack of basic knowledge of Myanmar’s ecosystem diversity, a lack of time-series of spatial data sufficient to estimate area change with confidence and poor accessibility across large areas of the country. Further work to improve time-series remote sensing models of ecosystems distributions is clearly a high priority, developed with temporal frequency sufficient to enable reliable projections across assessment timeframe (Murray et al. 2018; Lee et al. 2020). Data deficiency is therefore the greatest limitation to this study, and we particularly recommend improved networking among researchers and government departments to promote data sharing aimed at filling these substantial knowledge gaps. The new ecosystem typology for Myanmar developed for this study will help structure future studies of Myanmar’s ecosystems, while also assisting with ecosystem service assessment, identification of key biodiversity areas, and natural capital accounting (Bland et al. 2017; Murray et al. 2018).

### 4.3 Protecting Myanmar’s threatened ecosystems

The outlook for Myanmar’s ecosystems is at the crossroads. Comparisons with neighbouring countries of forest area and agricultural land as percentages of total land area reveal a clear pathway to sustaining a highly significant component of global biodiversity. According to World Bank syntheses, Myanmar ranks second highest in percent forest cover (43.6% forest) after Laos (82.1%), and followed by Thailand (32.2%), India (23.8%) and China (22.4%; The World Bank 2019). Thus, with more than 60% of the country covered by natural ecosystems, opportunities for conservation and restoration abound. Yet, with only 1.9% of the extent of Myanmar’s threatened ecosystems occurring within protected areas and recent increases in the rate of loss, swift action is required. This opens unique opportunities for strategic action to protect Myanmar’s ecosystems with a comprehensive, adequate and representative, well-managed protected area network and implementation of a range of ecosystem restoration actions.

Three challenges face authorities charged with improving the conservation status of Myanmar’s ecosystems. First, strategic expansion of protected areas (currently covering only 6.4% of Myanmar) is required to meet global targets and represent the full diversity of ecosystems and species. Priorities for expanding the protected area network should focus on conserving threatened ecosystems, focusing on regions where there is high spatial overlap of unprotected threatened intact ecosystems with rapidly emerging threats, and on biomes with high proportions of threatened component ecosystems (Table 2). Recent work proposing World Heritage status for a region in far north Myanmar is particularly promising, but highlights the need to engage with local communities to ensure successful environmental, social and economic outcomes while planning for biodiversity protection.

Second, despite expansive protected areas in northern Myanmar protecting large areas of intact wilderness, many local-scale threats continue to operate if protected areas are not managed effectively and co-operatively with local people. In this region, ecosystems currently assessed as Least Concern are continually at-risk of being uplisted into the threatened categories. Reported threats, even within protected areas, include poaching of characteristic species, loss of important structural elements due to high-value timber extraction, fragmentation due to roads and oil pipelines, and widespread degradation caused by unsustainable, short-term, agricultural practices (Bhagwat et al. 2017). This example illustrates that management of existing protected areas across Myanmar should continue to be improved, with appropriate resourcing, training and oversight, and regular reassessment of ecosystem status is necessary to identify ecosystems transitioning from lower categories of risk into the threatened categories.

Finally, overcoming aforementioned data deficiencies will be crucial to support efforts to improve the conservation of Myanmar’s natural ecosystems. Highest priorities include systematic biodiversity surveys and conducting targeted searches to confirm the continued existence of near-collapsed ecosystems, developing datasets to increase the number of red list criteria assessed per ecosystem, and continuing to build a deep knowledge base of Myanmar’s ecosystems through basic ecological research. It is worth noting the recent increase in local studies founded on detailed field observations (Oo & Lee 2007; Oo & Koike 2015; Khaing et al. 2019); it is critical to expand this work to other locations and build capacity by integrating these into national data inventories and global data stores such as the Global Biodiversity Information Facility.

## 5. Conclusion

Our study demonstrates the feasibility of conducting ecosystem risk assessments from a minimal information base, a situation experienced by many countries seeking to conserve their biological diversity. We developed the first comprehensive spatially explicit inventory of ecosystems for the country, applied simple time series and spatial analyses to represent responses to key pressures, classified ecosystems at different levels of risk and identified data deficient ecosystems in need of further investigation. We showed that nearly half of the 64 natural ecosystem types assessed met criteria for listing as threatened on the IUCN Red List of Ecosystems, emphasising the need for continued conservation and restoration action. The management of Myanmar’s natural ecosystems requires an integrated approach that continues to fill substantial knowledge gaps on the ecology, distribution and functioning of Myanmar’s ecosystems, while simultaneously implementing conservation actions to maintain, restore, and protect what remains.

## Acknowledgements

We thank the Global Environmental Facility and UNDP for providing funding for the Myanmar National Ecosystem Assessment under the “Strengthening Sustainability of Protected Areas Management in Myanmar” project (GEF #5159, UNDP #5162). In Myanmar, we thank His Excellency U Ohn Winn, Union Minister of the Ministry of Natural Resources and Environmental Conservation, Dr. Nyi Nyi Kyaw, Director General of the Forest Department and Dr. Naing Zaw Htun Director of the Nature and Wildlife Conservation Division for their strong support and interest in this program. We also thank U Saw Htun, Country Director of the WCS Myanmar Program for his advice and leadership during project implementation. We also thank the many Myanmar experts who participated in our expert meetings and guided the development and implementation of the project. This research was funded in part by grants from the US National Science Foundation (NSF 1457702) and the Helmsley Charitable Trust to KA. NM is the recipient of an Australian Research Council Australian Discovery Early Career Award (DE190100101) funded by the Australian Government. DK was supported by grants from the Australian Research Council (LP130100435, LP170101143). TW was supported by the International Climate Initiative (IKI) funded by The German Federal Ministry for the Environment, Nature Conservation and Nuclear Safety (BMU) on the basis of a decision adopted by the German Bundestag and by an anonymous gift to The Nature Conservancy.

## Appendices

**Table A.1.** Comprehensive list and summary statistics of Myanmar’s natural ecosystem types. Threatened ecosystems are identified by codes NT, Near Threatened; VU, vulnerable; EN, Endangered; CE, Critically Endangered; CO, Collapsed. Ecosystems with NA were not mapped during the ecosystem mapping process.

| Ecosystem Functional Group | Ecosystem Name | Global Ecosystem Code | Mapped area (km<sup>2</sup>) | Area protected (km<sup>2</sup>) | Proportion protected (%) | IUCN Status |
| --- | --- | --- | --- | --- | --- | --- |
| **REALM: TERRESTRIAL** |  |  |  |  |  |  |
| **BIOME: TROPICAL & SUBTROPICAL FORESTS** |  |  |  |  |  |  |
| Tropical/subtropical lowland rainforests | Tanintharyi island rainforests | MMR-T1.1.1 | 1936.67 | 183.57 | 9.48 | VU |
|  | Tanintharyi Sundaic lowland evergreen rainforest | MMR-T1.1.2 | 7165.65 | 9.65 | 0.13 | VU |
|  | Tanintharyi limestone tropical evergreen forest | MMR-T1.1.3 | 1935.94 | 0 | 0 | EN |
|  | Tanintharyi upland evergreen rainforest | MMR-T1.1.4 | 7719.44 | 21.14 | 0.27 | DD |
|  | Kayin evergreen tropical rainforest | MMR-T1.1.5 | 725.96 | 7.81 | 1.08 | EN |
|  | Southern Rakhine evergreen rainforest | MMR-T1.1.6 | 32.13 | 0 | 0 | CR |
|  | Western Shan Plateau subtropical evergreen rainforest | MMR-T1.1.7 | 7540.28 | 26.27 | 0.35 | VU |
|  | Kachin-Sagaing low elevation evergreen subtropical rainforest | MMR-T1.1.8 | 2699.54 | 1383.92 | 51.27 | DD |
|  | Kachin-Sagaing mid elevation subtropical rainforest | MMR-T1.1.9 | 4564.34 | 1862.77 | 40.81 | DD |
|  | Kachin hills subtropical rainforest | MMR-T1.1.10 | 7353.53 | 2531.81 | 34.43 | DD |
| Tropical/subtropical dry forests and scrubs | Tanintharyi semi-evergreen forest | MMR-T1.2.1 | 20502.64 | 1963.8 | 9.58 | DD |
|  | Rocky Tanintharyi karst | MMR-T1.2.2 | NA | NA | NA | DD |
|  | Mixed cane break | MMR-T1.2.3 | 248.97 | 4.23 | 1.7 | LC |
|  | Bago semi-evergreen forest | MMR-T1.2.4 | 7058.42 | 578.22 | 8.19 | CR |
|  | Dry zone foothills spiny scrub | MMR-T1.2.5 | 479 | 1.7 | 0.35 | DD |
|  | Rakhine hills bamboo brake | MMR-T1.2.6 | 7562.11 | 474.87 | 6.28 | LC |
|  | Rakhine hills semi-evergreen dry forest | MMR-T1.2.7 | 24884.06 | 1297.75 | 5.22 | VU |
|  | Magway dry cycad forest | MMR-T1.2.8 | 1107.8 | 18.99 | 1.71 | EN |
|  | Magway semi-evergreen dry gully forest | MMR-T1.2.9 | 2241.33 | 43.93 | 1.96 | VU |
|  | East Myanmar dry valley forest | MMR-T1.2.10 | 33946.63 | 53.08 | 0.16 | VU |
|  | Eastern Shan semi-evergreen forest | MMR-T1.2.11 | 28713.41 | 53.77 | 0.19 | VU |
|  | Western Shan semi-evergreen forest | MMR-T1.2.12 | 47563.58 | 418.35 | 0.88 | VU |
|  | Indaing forest | MMR-T1.2.13 | 39022.86 | 718.04 | 1.84 | EN |
|  | Northern semi-evergreen forest | MMR-T1.2.14 | 76735.19 | 12578.63 | 16.39 | DD |
| Tropical/subtropical moist montane rainforests | Tanintharyi cloud forest | MMR-T1.3.1 | 0.69 | 0.13 | 19.18 | CR |
| **BIOME: TEMPERATE-BOREAL FORESTS & WOODLANDS** |  |  |  |  |  |  |
| Boreal and temperate montane forests and woodlands | Kachin mountain conifer forest | MMR-T2.1.1 | 490 | 421.26 | 85.97 | DD |
| Warm temperate rainforests | Shan warm temperate rainforest | MMR-T2.4.1 | 7630.63 | 16.45 | 0.22 | EN |
|  | Chin Hills warm temperate rainforest | MMR-T2.4.2 | 11188 | 1464.47 | 13.09 | VU |
|  | Sagaing warm temperate rainforest | MMR-T2.4.3 | 5838.47 | 992.92 | 17.01 | LC |
|  | Kachin warm temperate rainforest | MMR-T2.4.4 | 11270.48 | 4922.49 | 43.68 | DD |
|  | Mountain bamboo brake | MMR-T2.4.5 | NA | NA | NA | DD |
|  | Kachin montane temperate broadleaf forest | MMR-T2.4.6 | 10542.3 | 4390.44 | 41.65 | DD |
| **BIOME: SAVANNAS AND GRASSLANDS** |  |  |  |  |  |  |
| Pyric tussock savannas | Rakhine coastal savanna | MMR-T4.2.1 | 5091.8 | 2.07 | 0.04 | DD |
|  | Central Ayeyarwady Than-Dahat grassy forest | MMR-T4.2.2 | 6838.38 | 76.48 | 1.12 | VU |
|  | Central Ayeyarwady Palm savanna | MMR-T4.2.3 | NA | NA | NA | CO |
|  | Shwe Settaw Sha-Bamboo thicket | MMR-T4.2.4 | 59.6 | 9.04 | 15.16 | NT |
|  | Magway Than-Dahat dry grassy forest | MMR-T4.2.5 | 4977.01 | 299.23 | 6.01 | LC |
|  | Sha thorny scrub | MMR-T4.2.6 | 652.8 | 1.25 | 0.19 | VU |
|  | Shan foothills Than-Dahat grassy forest | MMR-T4.2.7 | 6387.3 | 279.52 | 4.38 | VU |
|  | Shan hills pine savanna | MMR-T4.2.8 | 3566.91 | 19.19 | 0.54 | EN |
|  | Chin hills pine savanna | MMR-T4.2.9 | 1653 | 49.02 | 2.97 | LC |
|  | Sagaing hills pine savanna | MMR-T4.2.10 | 31.72 | 20.61 | 64.98 | LC |
|  | Kachin pine savanna | MMR-T4.2.11 | 9.5 | 1.8 | 18.95 | LC |
| Temperate grasslands | Shan plateau grasslands | MMR-T4.5.1 | NA | NA | NA | DD |
| **BIOME: POLAR/ALPINE** |  |  |  |  |  |  |
| Ice sheets, glaciers and perennial snowfields | Kachin snowfields | MMR-T6.1.1 | 3142.58 | 2237.59 | 71.2 | NT |
| Polar/alpine rocky outcrops | Alpine cliffs and screes | MMR-T6.2.1 | 296.34 | 292.46 | 98.69 | LC |
| Temperate alpine meadows and shrublands | High mountain scrub | MMR-T6.4.1 | 303.21 | 157.19 | 51.84 | LC |
|  | Alpine herbfield | MMR-T6.4.2 | 69.32 | 69.32 | 100 | EN |
| **REALM: SUBTERRANEAN** |  |  |  |  |  |  |
| **BIOME: DRY SUBTERRANEAN** |  |  |  |  |  |  |
| Subterranean lithic systems | Aerobic Karst caves | MMR-S1.1.1 | 0.38 | 0.01 | 1.99 | LC |
| **REALM: FRESHWATER/TERRESTRIAL** |  |  |  |  |  |  |
| **BIOME: PALUSTRINE WETLANDS** |  |  |  |  |  |  |
| Tropical flooded forests and peat forests | Ayeyarwady kanazo swamp forest | MMR-TF1.1.1 | NA | NA | NA | CR |
|  | Central dry evergreen riparian forest | MMR-TF1.1.2 | 87.11 | 0.02 | 0.02 | CR |
|  | Mixed delta scrub | MMR-TF1.1.3 | 825.13 | 0.25 | 0.03 | LC |
| Seasonal floodplain marshes | Ayeyarwady floodplain wetlands | MMR-TF1.4.1 | 811.25 | 0.05 | 0.01 | EN |
|  | Central Ayeyarwady floodplain grasslands | MMR-TF1.4.2 | 2333.8 | 0.51 | 0.02 | CR |
| **REALM: FRESHWATER** |  |  |  |  |  |  |
| **BIOME: LAKES** |  |  |  |  |  |  |
| Freeze-thaw freshwater lakes | Glacial lakes | MMR-F2.4.1 | 21.51 | 21 | 97.62 | LC |
| **REALM: MARINE/TERRESTRIAL** |  |  |  |  |  |  |
| **BIOME: SHORELINE SYSTEMS** |  |  |  |  |  |  |
| Muddy shores | Coastal mudflats | MMR-MT1.2.1 | 2997.68 | 6.92 | 0.23 | LC |
| Sandy shores | Sandy shoreline | MMR-MT1.3.1 | 1460.92 | 11.92 | 0.82 | LC |
| **BIOME: SUPRALITTORAL COASTAL SYSTEMS** |  |  |  |  |  |  |
| Coastal shrublands and grasslands | Tanintharyi coastal dune forest | MMR-MT2.1.1 | NA | NA | NA | DD |
|  | Rakhine coastal dune forest | MMR-MT2.1.2 | NA | NA | NA | DD |
| **REALM: MARINE/FRESHWATER/TERRESTRIAL** |  |  |  |  |  |  |
| **BIOME: BRACKISH TIDAL SYSTEMS** |  |  |  |  |  |  |
| Intertidal forests and shrublands | Tanintharyi mangrove forest | MMR-MFT1.2.1 | 3273.76 | 7.11 | 0.22 | NT |
|  | Ayeyarwady delta mangrove forest | MMR-MFT1.2.2 | 1235.23 | 126.68 | 10.26 | EN |
|  | Dwarf mangrove (shrubland) on shingle | MMR-MFT1.2.3 | 0.45 | 0 | 0 | CR |
|  | Rakhine mangrove forest on mud | MMR-MFT1.2.4 | 1801.41 | 0 | 0 | CR |
| Coastal saltmarshes | Grassy saltmarsh | MMR-MFT1.3.1 | NA | NA | NA | DD |

**Figure A.1.**
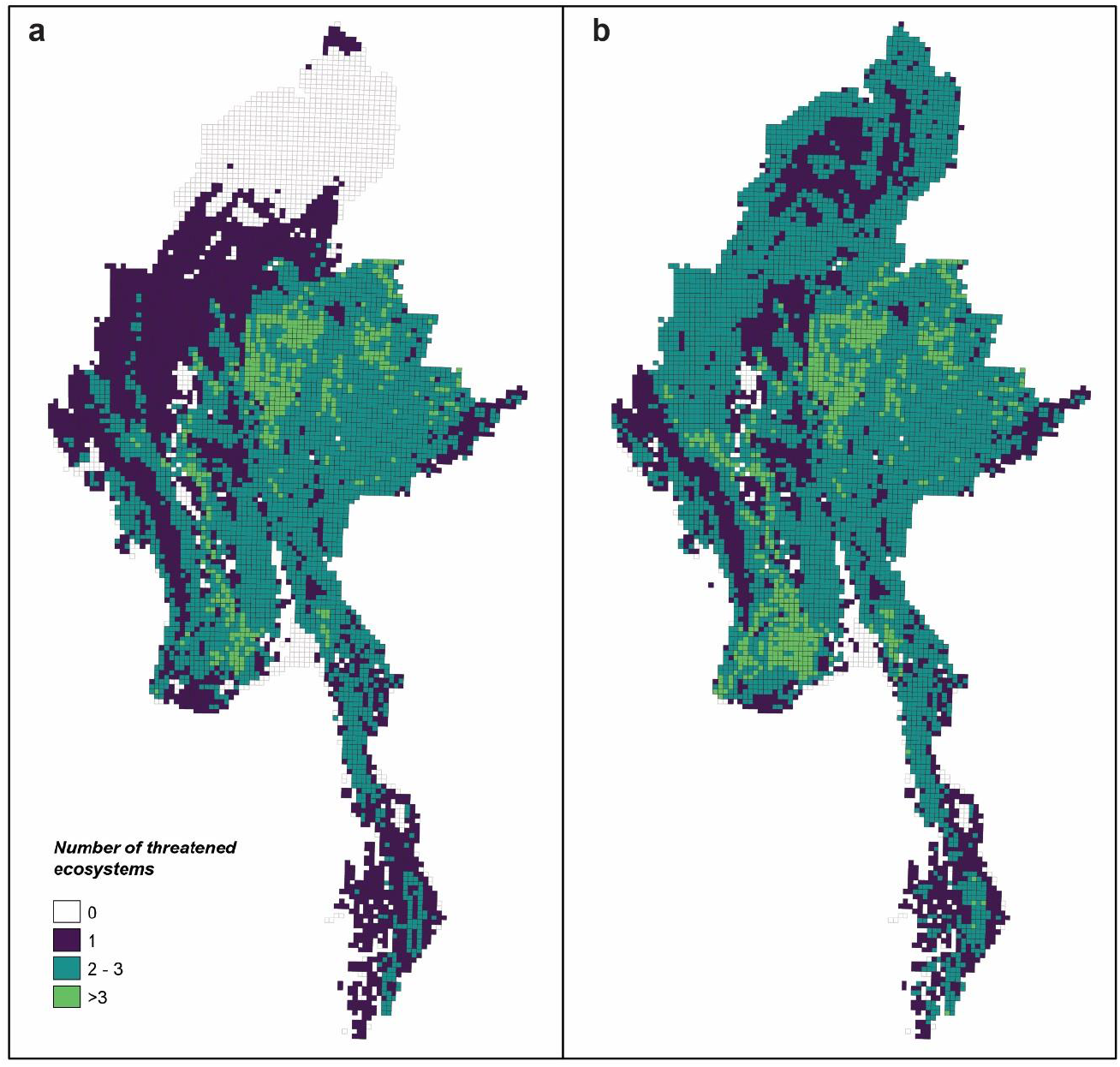
Uncertainty in the distribution of threatened ecosystems in Myanmar. The figure shows the “threatened ecosystem richness”, formulated as the sum of 10 × 10 area of occupancy cells in which an ecosystem considered threatened under the IUCN Red List of Ecosystems occurs. Data presented here are lower (a) and upper (b) plausible bounds reflect uncertainty in assessment outcomes as a result of lack of suitable data, model uncertainty, or expert judgement. Some white areas in central Myanmar are dominated by agriculture and therefore have no natural ecosystems (Refer to Figure 3).

